# Subcutaneous BCG vaccination protects against streptococcal pneumonia via regulating innate immune responses in the lung

**DOI:** 10.1101/2022.09.29.510220

**Authors:** Alisha Kang, Gluke Ye, Ramandeep Singh, Sam Afkhami, Jegarubee Bavananthasivam, Xiangqian Luo, Maryam Vaseghi-Shanjani, Anna Zganiacz, Mangalakumari Jeyanathan, Zhou Xing

## Abstract

Bacillus Calmette-Guérin (BCG) still remains the only licensed vaccine for TB and has been shown to provide nonspecific protection against unrelated pathogens. This has been attributed to the ability of BCG to modulate the innate immune system, known as trained innate immunity (TII). TII is associated with innate immune cells being in a hyper-responsive state leading to enhanced host defense against heterologous infections. Both epidemiological evidence and prospective studies demonstrate cutaneous BCG vaccine-induced TII provides enhanced innate protection against heterologous pathogens. Regardless of the extensive amounts of progress made thus far, the effect of cutaneous BCG vaccination against heterologous respiratory bacterial infections and the underlying mechanisms remain unknown. Here we show for the first time that s.c BCG vaccine-induced TII provides enhanced heterologous innate protection against pulmonary *S. pneumoniae* infection. We further demonstrate that this enhanced innate protection is mediated by accelerated neutrophilia in the lung and is independent of centrally trained circulating monocytes. New insight from this study will help design novel effective vaccination strategies against unrelated respiratory bacterial pathogens.

## Introduction

Bacillus Calmette-Guérin (BCG), the only licensed vaccine for tuberculosis (TB), remains the mostly widely administered vaccine worldwide given via the skin (1). Although it has limited efficacy in protection against pulmonary TB in adults, BCG vaccination has been found to provide nonspecific protective effects against unrelated heterologous infections beyond TB in humans (2–4). This has been attributed to the ability of live attenuated BCG to modulate the innate arm of the immune system and has been termed trained innate immunity (TII) (5). TII is associated with a state of hyper-responsiveness and leads to enhanced host defense (6).

The identification of beneficial nonspecific protective effects of BCG came from epidemiological evidence in which cutaneous BCG vaccination reduced all-cause mortality in low-birth-weight infants and the infants with a BCG scar had significantly lower mortality in early childhood (7, 8). In part, it is due to reduced risk of developing lower respiratory tract infections unrelated to TB (9, 10). Indeed, in prospective clinical studies, cutaneous BCG vaccination was found to protect healthy adults from yellow fever virus and malaria and reduced the incidence of respiratory infections in the elderly (11–14). BCG vaccine-mediated nonspecific innate immune protection is linked to its activating effects on circulating innate immune cells in the peripheral blood including monocytes, neutrophils, and NK cells associated with epigenetic and metabolic modifications and increased production of pro-inflammatory cytokines (15–18). Since most of the circulating innate immune cells such as monocytes are mature and thus have a short half-life, BCG vaccine-induced persisting changes in circulating innate immune cells has been found to result from its training effects on hematopoietic progenitor cells in the bone marrow (19, 20).

Preclinical studies have also demonstrated TII by parenteral or respiratory mucosal routes of BCG vaccination against heterologous pathogens including *Candida albicans, Streptococcus pneumoniae* and influenza virus or the target pathogen *M. tuberculosis* (21–23). Of note, however, most of the preclinical parenteral BCG studies have demonstrated the induction of TII only following intravenous (i.v) inoculation of BCG vaccine (17, 21, 24). While few studies have made a head-to-head comparison with cutaneous BCG vaccination, they conclude that different from i.v. or respiratory mucosal route of vaccination, cutaneous BCG vaccination is ineffective in TII induction (22, 23). Thus, up to date, these preclinical observations together have been at odds with human studies in that clinically or in real-world practice, BCG vaccine is always administered via the skin. Lack of clinically relevant preclinical models of cutaneous BCG vaccination and TII induction in the lung has hindered the mechanistic understanding of cutaneous BCG vaccine-induced TII in humans.

To fill the above knowledge gap, in the current study we have developed a murine model to study subcutaneous (s.c) BCG vaccine-induced TII in the lung. We show that s.c BCG vaccination with a small-dose inoculum induces TII in the lung which translates to enhanced innate immune protection against pulmonary *Streptococcus pneumoniae* (*S. pneumoniae*) infection. We provide evidence that s.c BCG vaccine-induced anti-streptococcal TII is mediated through accelerated neutrophilia in the lung. We further demonstrate that such enhanced protection is independent of centrally trained circulating monocytes. Our study sheds new light on the mechanisms via which cutaneous BCG vaccination offers TII against heterologous respiratory pathogens in mouse lungs.

## Results

### Enhanced innate immune protection against pulmonary *S. pneumoniae* infection in BCG vaccinated hosts

To begin addressing the subcutaneous (s.c.) BCG vaccine-induced TII in the lung, we adapted a murine model of *S. pneumoniae* respiratory infection. C57BL/6 mice vaccinated with s.c BCG for 5 weeks (wks) were infected intratracheally (i.t) with *S. pneumoniae* serotype 3 and evaluated for survival and clinical outcomes as a measure of weight loss and clinical signs (**Figure 1A**). A set of unvaccinated mice were also infected with *S. pneumoniae* as controls. Compared to control mice, a significantly greater number of BCG-vaccinated mice survived the infection (**Figure 1B**). While ∼70% of BCG vaccinated mice survived the infection, only 20% of control mice survived the infection. Of interest, BCG-vaccinated mice hardly lost any weight during the infection and exhibited minimal clinical signs. In stark contrast, a significant number of unvaccinated mice reached end point by 4d post-infection (∼50%) and exhibited severe clinical signs (**Figures 1C/D**). We next assessed the bacterial burden in the lung and spleen by colony forming units (CFU) assay (**Figure 1E**). Keeping in line with significantly improved clinical outcomes, BCG vaccinated mice better controlled bacterial replication both locally in the lung and systemically in the spleen (**Figure 1F**). BCG vaccinated mice carried ∼1.5 log less bacteria in the lung and spleen compared to their control counterparts. Of note, lung bacterial counts significantly increased over the period in control mice, in BCG vaccinated hosts bacterial replication was better controlled as indicated by reduced bacterial burden at 72h compared to 48h post-infection. We next evaluated the microscopic histological changes in the lung, s.c BCG vaccination did not cause any alterations to the lung, indicated by comparable histology to control counter parts (**Figure 1G**). Pulmonary *S. pneumoniae* infection led to marked changes in the lung of control animals as indicated by cell infiltration into the airways and around the bronchi which was accompanied by mild epithelial sloughing. In stark contrast, integrity of lung histology remained unaffected in BCG vaccinated hosts post-infection (72h). Next, we evaluated the longevity of BCG mediated protection against pulmonary *S. pneumoniae* infection (**Figure S1A)**. C57BL/6 mice vaccinated with s.c BCG for 16wks or left unvaccinated were infected i.t with *S. pneumoniae* and clinical outcomes and bacterial burden were assessed. BCG vaccinated hosts were modestly protected against *S. pneumoniae* as indicated by improved clinical outcomes (**Figures S1B/C**) accompanied by lower bacterial burden in the lung and spleen compared to control animals (**Figure S1D**). The above data indicate that parenteral BCG vaccination leads to heterologous protection against pulmonary *S. pneumoniae* infection.

**Figure 1.**
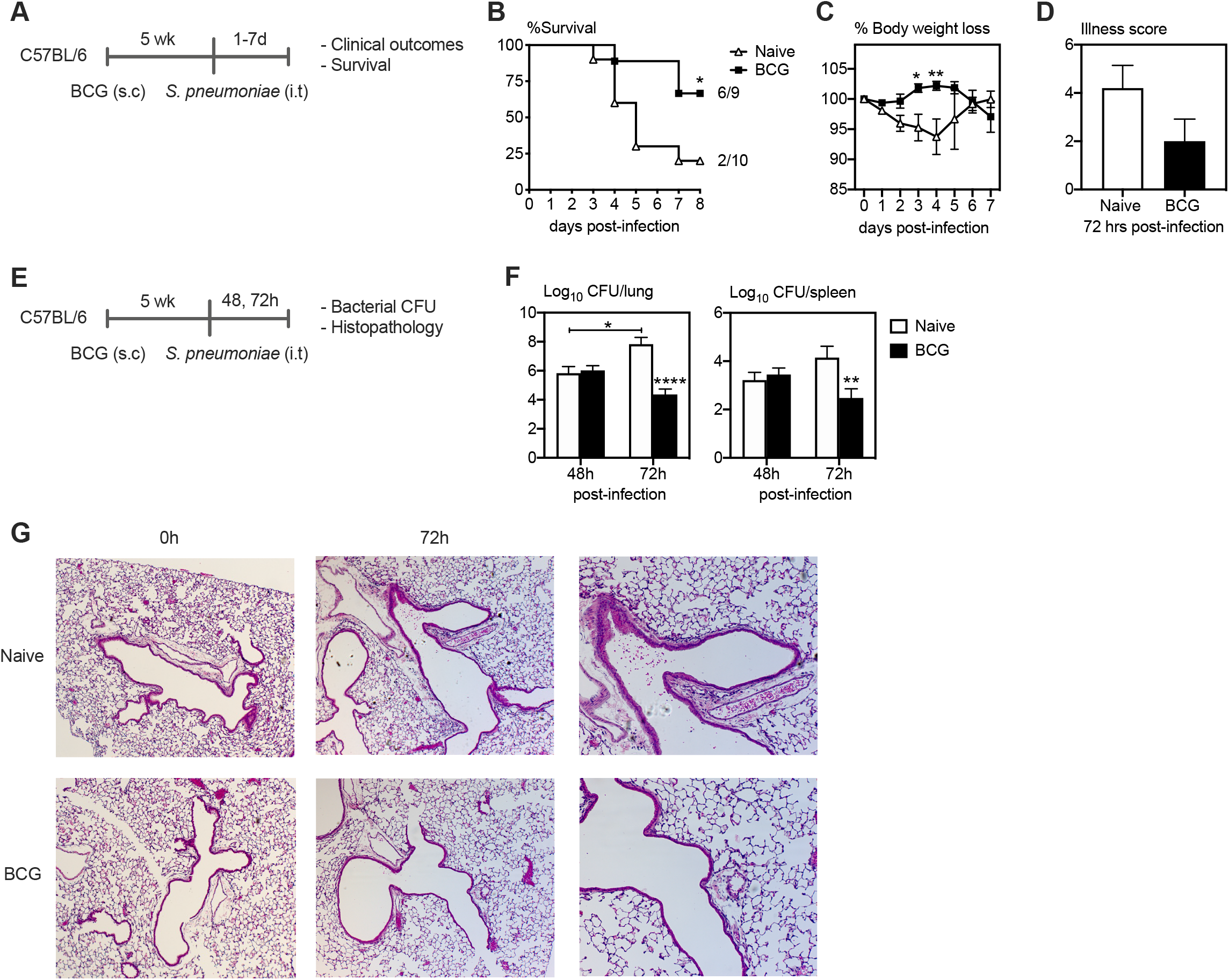
Enhanced protection against pulmonary *S. pneumoniae* infection in BCG vaccinated hosts. A) Experimental schema. B) Survival of BCG vaccinated mice post-*S. pneumoniae* infection. C & D) Changes in body weight and illness score one-week post-*S. pneumoniae* infection. E) Experimental schema. F) Bacterial counts (CFU) in the lung and spleen at 48 and 72h post-*S. pneumoniae* infection in control and BCG vaccinated mice. G) Lung histopathological images at 0 and 72h post-*S. pneumoniae* infection in control and BCG vaccinated mice. Data presented in (B, C, D and F) represent mean ±SEM. Data are representative of 1 – 2 independent experiments, n = 5 – 10 mice/group. Statistical analysis for (B, C and F) were two-way ANOVA with Sidak’s multiple comparisons test, (D) was two-tailed unpaired t-test. *p < 0.05; **p < 0.01; ***p < 0.001; ****p < 0.0001.

### Accelerated and enhanced neutrophilia in the lung following pulmonary *S. pneumoniae* infection in BCG vaccinated hosts

We next evaluated whether enhanced protection in BCG vaccinated host was accompanied by differential cellular and molecular immune responses in the lung following pulmonary *S. pneumoniae* infection. To this end, cellular responses were profiled before (0h) and at 18/36h post-*S. pneumoniae* infection in the airways and lung of BCG vaccinated and control hosts (**Figure 2A**). Airway cells isolated by bronchoalveolar lavage (BAL) and lung mononuclear cells were immunostained with a comprehensive panel of innate immune surface markers as described previously and analyzed by flow cytometry (25). The airways were predominately comprised of alveolar macrophages (AM) both in control and BCG vaccinated hosts before *S. pneumoniae* infection (0h) (data not shown). Upon infection, the total number of neutrophils were comparable between the two groups at 18h but markedly increased in BCG vaccinated hosts at 36h post-infection (**Figures 2B/C**). Of interest, neutrophils were not recruited to the airways until 36h post-infection in control mice. The total number of alveolar and interstitial macrophages (IMs) increased post-infection in both groups with marginally increased numbers in BCG vaccinated hosts at 36h post-infection (**Figure 2D**). In comparison, cellular responses in the lung did not differ between groups but an increase in cellular infiltration was observed at 36h post-infection (**Figure E-H**).

**Figure 2.**
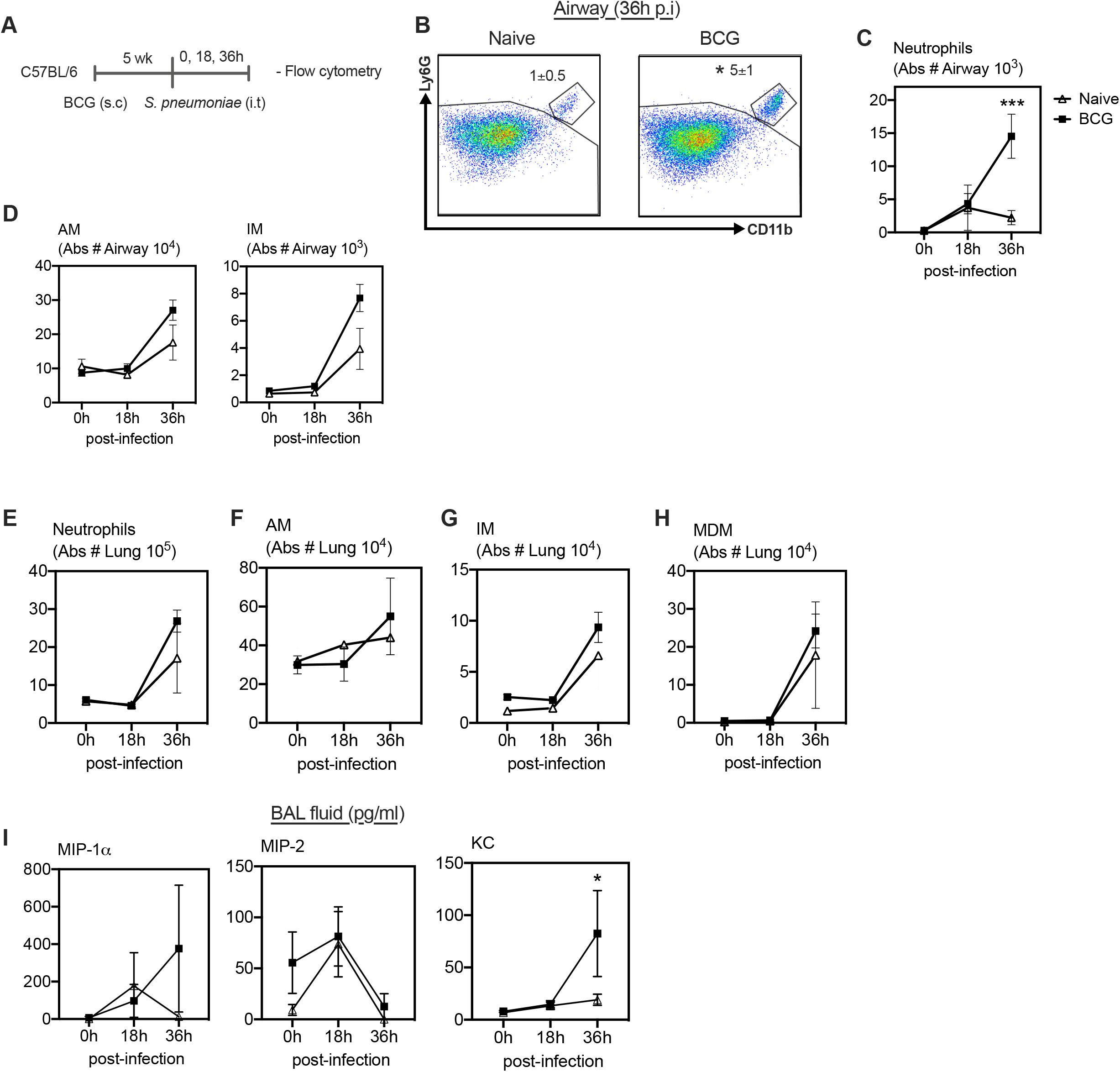
Accelerated and enhanced neutrophilia in the lung following pulmonary *S. pneumoniae* infection in BCG vaccinated hosts. A) Experimental schema. B) Dot plots of gated mononuclear cells from the BAL of control and BCG vaccinated mice. Analysis was performed utilizing default FlowJo V.10 software settings. C & D) Absolute cell counts in BAL at 18 and 36h post-*S. pneumoniae* infection. E-H) Absolute cell counts in the lung at 18 and 36h post-*S. pneumoniae* infection. I) Levels of neutrophilic chemokines in the BAL at 18 and 36h post-*S. pneumoniae* infection. Data presented in (B – I) represent mean ±SEM. Data are representative of 1 experiment, n = 4 mice/group. Statistical analysis for (C – I) were two-way ANOVA with Sidak’s multiple comparisons test. *p < 0.05; **p < 0.01; ***p < 0.001; ****p < 0.0001.

Given the accelerated and enhanced neutrophilia in the airways, we then examined neutrophil attracting chemokine protein levels in the BAL fluids at various time points post-infection. Before infection (0h), the levels of KC, MIP-2 and MIP-1α were low in the airways of both control and BCG vaccinated hosts. Following infection, these chemokine levels were largely comparable between the two groups at 18h post-infection (**Figure 2J**). However, keeping in line with neutrophil infiltration to the airways at 36h post-infection (**Figure 2C**), neutrophil attracting chemokine, KC protein levels were significantly increased in the airways of BCG vaccinated hosts at 36h post-infection (**Figure 2J**). The above data suggest that there is an association between accelerated and enhanced neutrophilia in the lung and improved protection against pulmonary *S. pneumoniae* infection in BCG vaccinated hosts.

### Heightened bactericidal activity and improved protection against *S. pneumoniae* infection mediated by alveolar macrophages from BCG vaccinated hosts

Since alveolar macrophages are part of the first line of defence against respiratory tract infections and have been shown to play a vital role in protection against *S. pneumoniae* infection (26), we next evaluated the protective role of alveolar macrophages from BCG vaccinated hosts. To this end, AMs harvested from the airways of control (Naïve-AM) and BCG vaccinated mice (BCG-AM) were infected *ex vivo* with *S. pneumoniae* (**Figure 3A**). Phagocytic capacity and killing rates were determined at 1 and 2h post-*ex vivo* infection, respectively, by CFU assay. Compared to Naive-AMs, BCG-AMs demonstrated marginally increased phagocytic capacity as indicated by ∼2000 more intracellular bacteria in BCG-AMs (**Figure 3B**) and markedly enhanced killing capacity (**Figure 3C**). Since AMs from BCG vaccinated mice exhibited enhanced bactericidal activity, we wanted to further assess the role of these AMs in enhanced protection against *S. pneumoniae* infection *in vivo*. AMs harvested from the airways of control and BCG vaccinated donor mice were adoptively transferred to naïve mice. At 4h after transferring AMs, these mice were infected with *S. pneumoniae* and clinical outcomes and bacterial burden were assessed (**Figure 3D**). BCG-AM transferred mice demonstrated significantly improved protection compared to those transferred with Naïve-AM by sustaining their body weight and without showing any clinical signs of disease (**Figures 3E/F**). Although bacterial burden in the lung and spleen were comparable between the two groups at 72h post-infection (**Figure 3G**), BCG-AM transferred lungs demonstrated limited histological changes compared to the lung of Naïve-AM transferred mice, which displayed diffused alveolar hemorrhages and infiltration of immune cells within the airways (**Figure 3H**). These data together indicate, with the enhanced protection observed in BCG vaccinated hosts, that parenteral BCG vaccination imparted changes in resident alveolar macrophages leads to induction of TII in the lung against heterologous *S. pneumoniae* infection.

**Figure 3.**
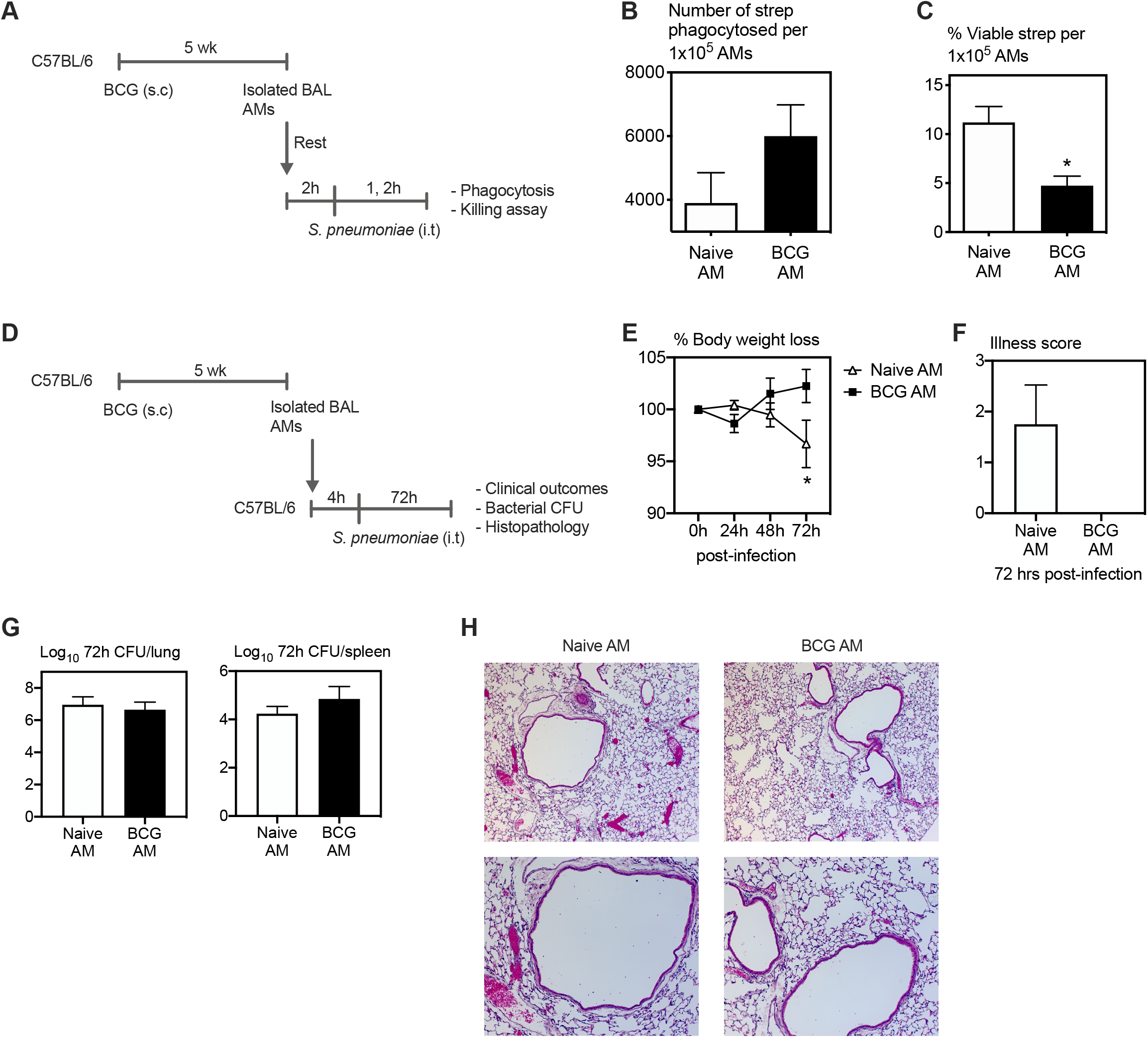
Heightened bactericidal activity and improved protection against *S. pneumoniae* infection mediated by AMs from BCG vaccinated hosts. A) Experimental schema. B) Phagocytosis as intracellular CFU of *S. pneumoniae* by AMs isolated from control and BCG vaccinated mice. C) Killing rates of phagocytosed *S. pneumoniae* by AMs at 1 and 2h post-*ex vivo S. pneumoniae* infection. D) Experimental schema. E & F) Changes in body weight and illness score post-*S. pneumoniae* infection. G) Bacterial counts (CFU) in the lung and spleen at 72h post-*S. pneumoniae* infection in control and BCG vaccinated mice. H) Lung histopathological images at 72h post-*S. pneumoniae* infection in control AM transferred and BCG AM transferred mice. Data presented in (B, C, E – G) represent mean ±SEM. Data are representative of 1 – 2 independent experiments, n = 3 – 5 mice/group. Statistical analysis for (B, C, F and G) were two-tailed unpaired t-tests, for (E) was two-way ANOVA with Sidak’s multiple comparisons test. *p < 0.05; **p < 0.01; ***p < 0.001; ****p < 0.0001.

### Neutrophilia in the lung during the early stages of infection plays a protective role against *S. pneumoniae* infection in BCG vaccinated hosts

Given that enhanced protection against *S. pneumoniae* infection in BCG vaccinated hosts was accompanied by enhanced and accelerated recruitment of neutrophils to the respiratory mucosa, we next evaluated the direct role of neutrophils in such enhanced protection. To this end, using a neutrophil depleting mAb (αLy6G) we partially depleted neutrophils *in vivo* immediately after *S. pneumoniae* infection in BCG vaccinated mice (BCG α-Ly6G) as previously described (27) (**Figure 4A**). As a control group, BCG vaccinated mice were administered isotype antibodies (BCG isotype). Mice were monitored for changes in weight, and clinical signs for 72h post-infection. A set of mice were sacrificed at either 48 or 72h post-infection and bacterial burden in the lung and spleen were assessed. Compared to isotype controls, the mice partially depleted of neutrophils (BCG α-Ly6G) displayed markedly increased weight loss and clinical signs of disease (**Figures 4B/C**). This was accompanied by significantly increased bacterial load in the lung at both 48 and 72h post-infection (**Figure 4D**). Of interest, in the presence of intact neutrophil responses (BCG isotype), bacterial burden in the lung declined significantly at 72h compared to 48h, while it remained unchanged in animals partially depleted of neutrophils (BCG α-Ly6G). A similar trend was seen in the spleen, where at 72h post-infection bacteria was undetectable (**Figure 4D**). These data suggest that accelerated neutrophilia in BCG vaccinated mice plays a critical role in enhanced heterologous innate protection against *S. pneumoniae* infection.

**Figure 4.**
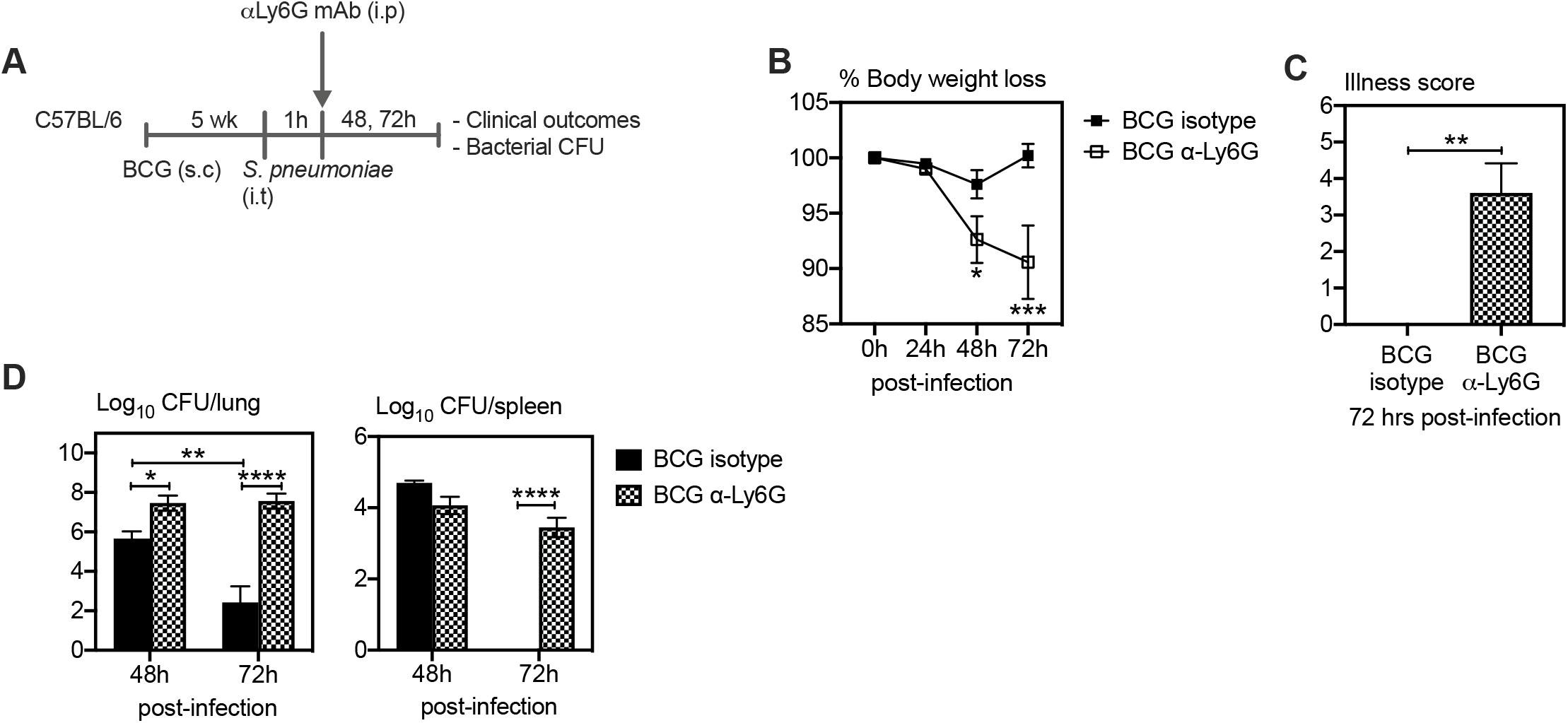
Accelerated neutrophilia plays a critical role in protection against *S. pneumoniae* infection in BCG vaccinated hosts. A) Experimental schema. B & C) Changes in body weight and illness score post-*S. pneumoniae* infection. D) Bacterial counts (CFU) in the lung and spleen at 48 and 72h post-*S. pneumoniae* infection in control and BCG vaccinated mice. Data presented in (B – D) represent mean ±SEM. Data are representative of 2 independent experiments, n = 5 mice/group. Statistical analysis for (B and D) were two-way ANOVA with Sidak’s multiple comparisons test, (C) was two-tailed unpaired t-test. *p < 0.05; **p < 0.01; ***p < 0.001; ****p < 0.0001.

### Enhanced protection against *S. pneumoniae* infection in BCG vaccinated hosts is independent of monocytes

Given that i.v BCG vaccine-induced heterologous protection has been attributed to BCG induced trained innate immunity in monocytes (20, 21) we next evaluated the relative role of monocytes in protection against *S. pneumoniae* infection in cutaneous BCG vaccinated hosts. To address this, CCR2^-/-^ mice lacking classical Ly6C^hi^ monocytes (CCR2^-/-^ BCG) or WT mice (WT BCG) were BCG-vaccinated for 5 wks. Groups of WT (WT naïve) or CCR2^-/-^ (CCR2^-/-^ naïve) mice were left unvaccinated as controls. Mice in all groups were infected with *S. pneumoniae* (**Figure 5A**). Mice were monitored daily for clinical signs and weight loss and bacterial burden in the lung and spleen were assessed by bacterial CFU assay at 48h post infection, by which time ∼70% of unvaccinated CCR2^-/-^ mice reached endpoint. Compared to unvaccinated WT mice (WT naïve), unvaccinated CCR2^-/-^ (CCR2^-/-^ naïve) mice displayed moderately increased weight loss and clinical signs (**Figures 5B/C**). In contrast, BCG vaccinated WT and CCR2^-/-^ mice maintained their body weight. In keeping with clinical outcomes, lungs of unvaccinated CCR2^-/-^ mice carried significantly increased number of bacteria compared to unvaccinated WT counterparts. In sharp contrast, comparable levels of bacterial burden were observed in BCG vaccinated WT and CCR2^-/-^mice (**Figure 5D**). A similar trend in bacterial burden was observed in the spleen (**Figure 5E**). The above data suggest that although monocytes play a critical role in protection against *S. pneumoniae* infection in unvaccinated hosts, it was dispensable in BCG vaccinated hosts.

**Figure 5.**
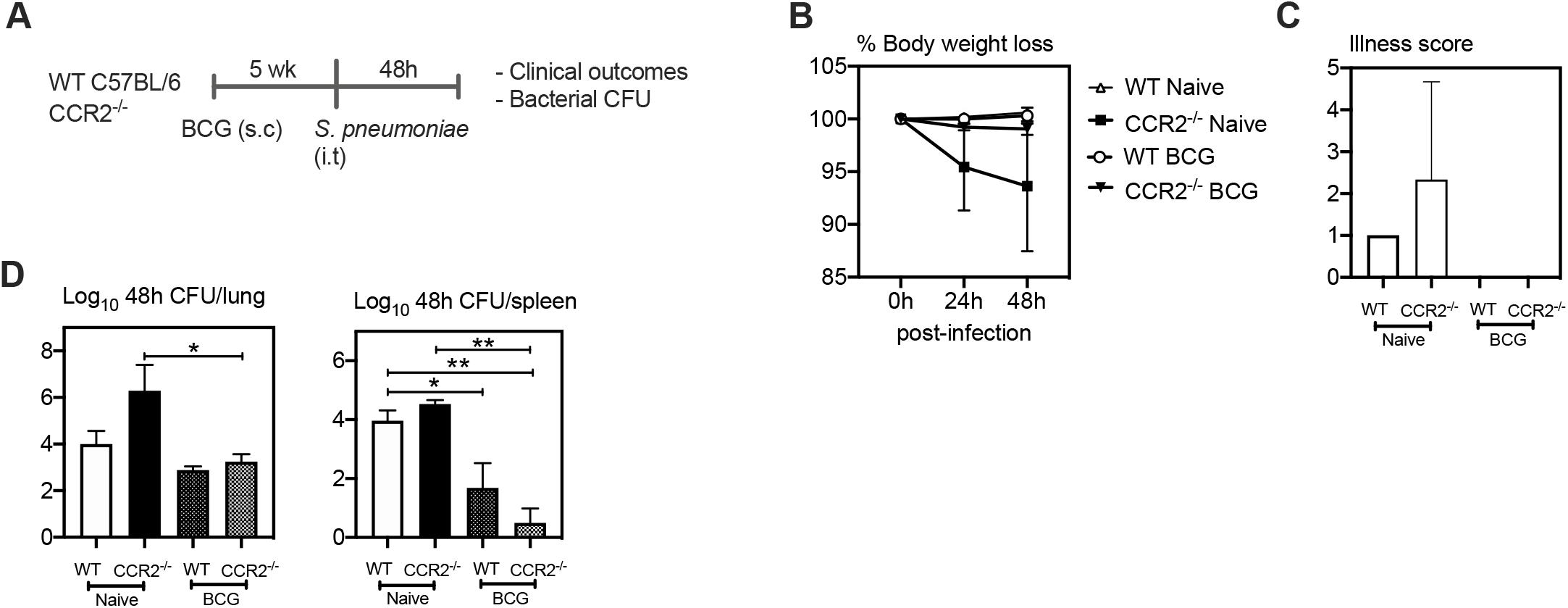
Enhanced protection against *S. pneumoniae* infection in BCG vaccinated hosts is independent of monocytes. A) Experimental schema. B & C) Changes in body weight and illness score post-*S. pneumoniae* infection. D) Bacterial counts (CFU) in the lung and spleen at 48h post-*S. pneumoniae* infection in control and BCG vaccinated mice. Data presented in (B – D) represent mean ±SEM. Data are representative of 1 experiment, n = 3 mice/group. Statistical analysis for (C) was two-way ANOVA with Sidak’s multiple comparisons test, (D) was one-way ANOVA with Tukey’s multiple comparisons test. *p < 0.05; **p < 0.01; ***p < 0.001; ****p < 0.0001.

## Discussion

Over the years, BCG vaccination has been shown to provide nonspecific protection against heterologous infections other than TB in humans (4–6). Mounting preclinical evidence attribute such protection to BCG vaccine-induced TII in monocytes following i.v BCG vaccination (17, 21, 24). These studies also suggest that compared to i.v BCG vaccination, cutaneous BCG vaccination is ineffective in TII induction (22, 23). Thus, up to date, these preclinical observations together have been at odds with human studies in that clinically or in real-world practice, BCG vaccine is always administered via the skin. Lack of clinically relevant preclinical models of cutaneous BCG vaccination and TII induction in the lung has hindered the mechanistic understanding of cutaneous BCG vaccine-induced TII in humans. To fill the above knowledge gap, in the current study we have developed a murine model to study s.c BCG vaccine-induced TII in the lung. Our study shows for the first time that s.c BCG vaccine-induced TII in the lung provides enhanced heterologous innate immune protection against pulmonary *S. pneumoniae* infection. Such enhanced protection was mediated via accelerated neutrophilia in the lung and independent of centrally trained circulating monocytes. Thus, our study reveals the mechanisms by which cutaneous BCG vaccination induces TII against heterologous respiratory bacterial infections in the lung and hold implications in developing strategies to modulate immunity at a remote tissue site.

To date, preclinical studies demonstrate that only i.v BCG vaccination is associated with TII and suggests that cutaneous BCG vaccination fails to induce TII. However, our findings that resident alveolar macrophages in BCG vaccinated hosts display heightened phagocytic and killing capacities indicate that cutaneous BCG vaccination can also remotely impact the functionality of resident AMs. This observation is further supported by a recent study that showed cutaneous BCG vaccination in mice modifies the phenotype of alveolar macrophages in the airways and alters the *in vivo* alveolar macrophage responses to *M. tb* infection (28). An ongoing study in our laboratory also demonstrated that s.c BCG vaccination induces memory AMs with the characteristics of TII which goes on to provide enhanced early protection against *M. tb* infection (unpublished). The current study provides further evidence that such TII in the lung following cutaneous BCG vaccination can also mediate heterologous protection against pulmonary *S. pneumoniae* infection. Interestingly, despite phenotypic and functional alterations in alveolar macrophages, gross cellularity in the airways is unaffected following cutaneous BCG vaccination, indicating that induction of TII in alveolar macrophages is independent of centrally trained monocytes (28, 29). Of note, although pulmonary *S. pneumoniae* infection led to the recruitment of centrally trained monocytes to the lung in BCG vaccinated hosts, they did not contribute to the enhanced protection as indicated by comparable clinical outcomes and bacterial control in BCG vaccinated CCR2^-/-^ and WT mice.

Although other studies have demonstrated that BCG vaccination provides nonspecific protection against heterologous viral, fungal, and bacterial pathogens we have shown that cutaneous BCG vaccination provides enhanced protection against pulmonary *S. pneumoniae* infection (19, 22, 30). Although much is known regarding the mechanisms of i.v BCG vaccine-induced TII and nonspecific protection, more preclinical mechanistic studies are needed to better understand cutaneous BCG vaccine-induced TII in humans. Thus far, BCG-mediated nonspecific innate protection has been mainly attributed to circulating monocytes, NK cells and neutrophils and is associated with epigenetic and metabolic reprogramming leading to increased production of pro-inflammatory cytokines (15, 16, 18, 31). It has been shown that parenteral BCG vaccination gains access to the bone marrow resulting in its training effects on hematopoietic progenitor cells (19, 32). Circulating innate immune cells, specifically neutrophils have been shown to play a critical role in host defense against bacterial infections including *S. pneumoniae* (33). Our findings suggest that cutaneous BCG vaccine-induced anti-streptococcal TII is mediated via accelerated neutrophilia in the lung. We show that partial depletion of neutrophils in BCG vaccinated mice significantly reduced protection. This is in line with our previous work in which TII induced by an adenoviral vectored vaccine augmented anti-streptococcal protection via accelerated neutrophilia in the lung (25). This demonstrates that cutaneous BCG vaccine-induced TII and enhanced innate protection against pulmonary *S. pneumoniae* infection is mediated by accelerated neutrophilia in the lung.

In conclusion, our study reveals that cutaneous BCG vaccine-induced TII in the lung can improve protection against pulmonary *S. pneumoniae* infection by accelerated and enhanced neutrophilia in the lung and such enhanced protection is independent of centrally trained monocytes. Such knowledge shall help design novel effective universal vaccination strategies against unrelated respiratory bacterial pathogens.

## Acknowledgments

This work was supported by the Canadian Institutes of Health Research (CIHR) Foundation Program, National Sanitarium Association of Canada, Canadian Foundation of Innovation.

## Author Contributions

AK, MJ and ZX conceived and designed the study. AK, GY, RS, SA, JB, XL, MV-S and AZ performed experiments. AK and ZX wrote the paper.

## Supplemental figure titles and legends

**Figure S1.**
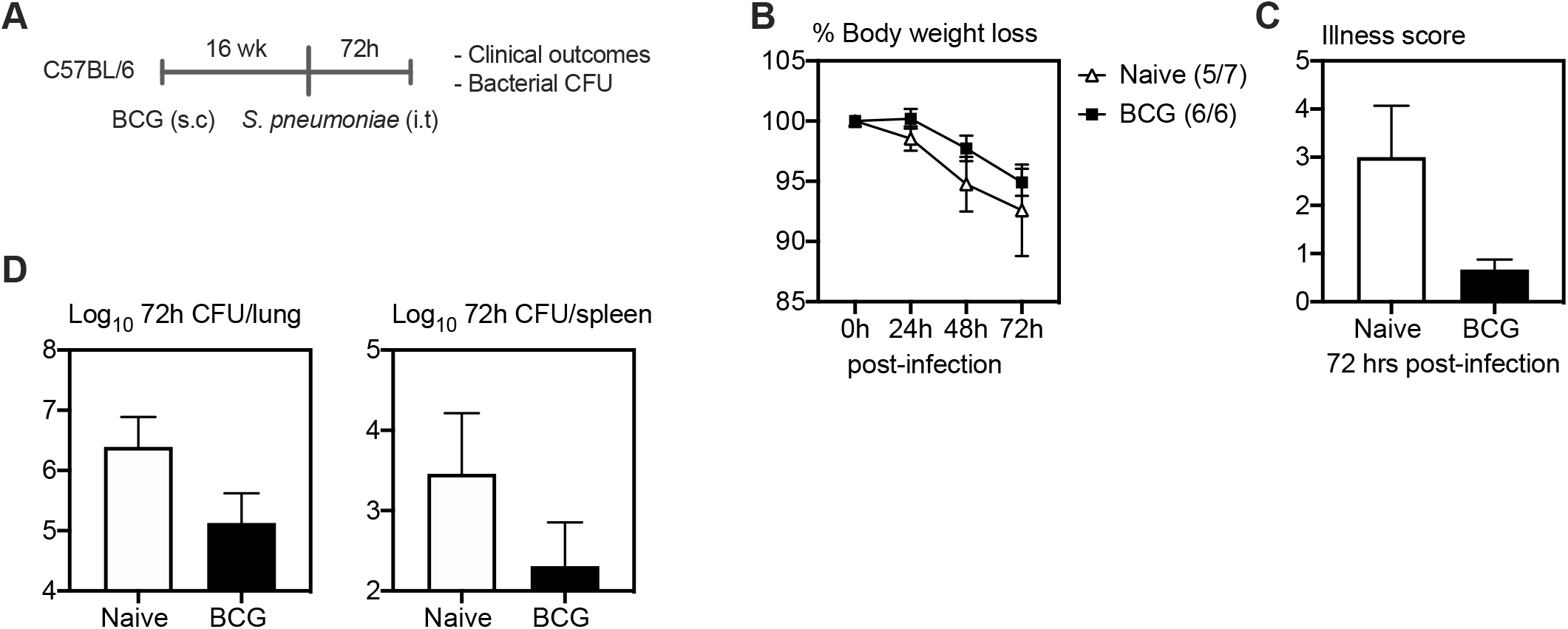
Enhanced protection against *S. pneumoniae* infection in BCG vaccinated hosts is long lasting. A) Experimental schema. B & C) Changes in body weight and illness score post-*S. pneumoniae* infection. D) Bacterial counts (CFU) in the lung and spleen at 72h post-*S. pneumoniae* infection in control and BCG vaccinated mice. Data presented in (B – D) represent mean ±SEM. Data are representative of 1 experiment, n = 6 – 7 mice/group. Statistical analysis for (B) was two-way ANOVA with Sidak’s multiple comparisons test, (C and D) were two-tailed unpaired t-tests. *p < 0.05; **p < 0.01; ***p < 0.001; ****p < 0.0001.

## Materials and methods

### Mice

Wild type 6–8-week-old female C57BL/6 mice were purchased from Charles River. Chemokine (C-C motif) receptor 2 (CCR2) knockout mice on a C57BL/6 background (B6.129S4-*Ccr2tm1Ifc*/J) were purchased from the Jackson Laboratory. All mice were housed in a specific pathogen free level B facility at McMaster University. All experiments were conducted in accordance with the guidelines of the animal research ethics board of McMaster University.

### Mycobacterial preparation for immunization

*M. bovis* BCG (Pasteur strain) was grown in Middlebrook 7H9 broth supplemented with Middlebrook oleic acid-albumin-dextrose-catalase (OADC) enrichment (Invitrogen Life Technologies, Carlsbad, CA), 0.002% glycerol, 0.05% Tween 80 for < 10-15 days, then aliquoted and stored in −70°C until needed. Prior to each use, bacilli were washed twice with phosphate-buffered saline (PBS) containing 0.05% Tween 80 and passed through a 27-gauge needle 10 times to disperse clumps. Mice were vaccinated subcutaneously (s.c) with 20 – 50,000 CFU/mouse of BCG in 100ul of PBS. The exact dose of vaccination was verified by tittering the inoculum using colony forming unit (CFU) assay.

### *Streptococcus pneumoniae* preparation for infection

A clinical isolate of *S. pneumoniae* serotype 3 (ATCC 6303; ATCC, Manassas, VA) was prepared. Briefly, the frozen bacterial stock was thawed in 37°C water bath and cultured at 37°C in 5% CO_2_ in Todd Hewitt broth (BD Biosciences, San Jose, CA) to mid-logarithmic phase, OD = 0.40 – 0.50. Bacteria were harvested and resuspended in cold PBS. For intratracheal (i.t) immunization, mice were infected with 10-50,000 CFU/mouse of *S. pneumoniae* in 40ul of PBS. The stock titer and infectious dose was verified by plating 10-fold serial dilutions on blood tryptic soy agar (BD Biosciences, San Jose, CA) supplemented with 5% defibrinated sheep blood (Hemostat, Dixon, CA) and 10 ug/ml neomycin (Sigma-Aldrich, St. Louis, MO).

### Bronchoalveolar lavage and lung mononuclear cell isolation

Mice were anesthetized by isoflurane and euthanized by exsanguination 5 weeks post-vaccination. Cells from bronchoalveolar lavage and lung tissue were isolated as previously described (34). Briefly, the lung was lavaged five times to a volume of 1.35 ml of PBS by cannulating the trachea using polyethylene tubing attached to 23-gauge needle and syringe. Following exhaustive bronchoalveolar lavage, lungs were cut into small pieces and digested in RPMI solution containing collagenase type 1 (ThermoFisher Scientific Waltham, MA) for 1 hr at 37°C in an agitating incubator. Single-cell suspension was obtained by crushing the digested lung tissue through a 100 μm basket filter (BD Biosciences, San Jose, CA) and subsequent lysis of RBCs by resuspending the cell pellet in Ammonium-Chloride-Potassium (ACK) lysing buffer for 2 min. Isolated cells were then resuspended in complete RPMI 1640 medium (RPMI 1640 supplemented with 10% FBS, 1% L-glutamine, 10 mM HEPES, 0.5mM Na pyruvate, 55 μM 2-Mercaptoethanol, 0.1mM NEAA, and with 1% penicillin/streptomycin) for flow cytometry staining. Isolated AMs from bronchoalveolar lavage were resuspended in PBS for adoptive transfer, mice were given 120,000 cells i.t in 40ul PBS.

### Immunostaining for cell phenotype characterization using flow cytometry

Cell immunostaining and flow cytometry were performed as previously described (35). Briefly, mononuclear cells from bronchoalveolar lavage (BAL) and the lung were plated in a U-bottom 96-well plate at a concentration of 20 million cells/ml in PBS. Following staining with Aqua dead cell staining kit (ThermoFisher Scientific Waltham, MA) at room temperature for 20-30 min, cells were washed and blocked to avoid non-specific staining with anti-CD16/CD32 (clone 2.4G2) in 0.5% BSA-PBS (FACS buffer) for 15 min on ice and then stained with fluorochrome-labeled mAbs for 20-30 min on ice [60]. Fluorochrome-labeled mAbs used for staining myeloid cells including alveolar macrophages, interstitial macrophages, monocyte-derived macrophages, monocytes, dendritic cells and neutrophils were anti-CD45-APC-Cy7 (clone 30-F11), anti-CD11b-PE-Cy7 (clone M1/70), anti-CD11c-APC (clone HL3), anti-MHC II-Alexa Flour 700 (clone M5/114.15.2), anti-CD3-V450 (clone 17A2), anti-CD45R-V450 (clone RA3-6B2), anti-Ly6C-Biotin (clone HK1.4), Streptavidin-Qdot800, anti-CD24-BV650 (clone M1/69), anti-CD64-PE (clone X54-5/7.1), anti-Ly6G-BV605 (clone 1A8), anti-Siglec-F-PE-CF594 (clone E50-2440), anti-CD80-PerCP-Cy5.5 (clone 16-10A1), anti-CD86-V450 (clone GL1), anti-CD284(TLR4)-APC (clone SA15-21), anti-CD11c-BV711 (clone HL3), anti-CD282(TLR2)-BV421 (clone CB225), anti-CD192(CCR2)-BV421 (clone SA203G11), anti-CD45.2-PerCP-Cy5.5 (clone 104). Immediately upon completion of the staining procedure, immunostained cells were processed according to the BD Biosciences instruction for flow cytometry and ran on a BD LSRII or BD LSRFortessa flow cytometer. Data were analyzed using FlowJo software (Version 10; Tree Star, Ashland, OR).

### Bacterial enumeration following *S. pneumoniae* challenge

The level of bacterial CFU in the lung, spleen, BALF and BAL cells was enumerated at various timepoints post-infection by plating neat and serial dilutions of tissue homogenates onto blood agar plates. Plates were dried, inverted, and placed into 5% CO_2_ incubator at 37°C for 12 – 16 hrs. Colonies were then enumerated and expressed as log_10_ CFU per organ.

### *Ex vivo* bacterial stimulation, phagocytosis and killing assays

Isolated alveolar macrophages from bronchoalveolar lavage were resuspended in complete RPMI medium without penicillin/ streptomycin (P/S-free media) and plated at 1.5×10^5^ cells/well in 48-well plates (100ul/well). Cells were incubated in a 37°C 5% CO_2_ cell-culture incubator for 1 hr and washed twice with P/S-free media. *S. pneumoniae* was grown as described above. Bacterial suspension was supplemented with 10% mouse serum (made in house from naïve and BCG C57BL/6 mice) and incubated for 30 min at 37°C and resuspended in P/S-free media. Bacteria were supplemented to cell culture wells at a MOI = 2.5 (3.75×10^5^ CFU of bacteria into 1.5×10^5^ cells/well). Cells were then incubated for 1 hr at 37°C, followed by supplementing with 500ul complete RPMI 1640 (containing 1% penicillin/streptomycin)/well and incubated for additional 30 min to remove any cell bound bacteria. Bacterial-stimulated alveolar macrophages were lysed at 1, 2, or 12 hrs post-removal of extracellular bacteria by adding 1ml per well of autoclaved MiliQ water. Cell lysates were diluted by 1:10 serial dilutions in PBS, plated on blood agar and cultured overnight. Bacterial phagocytosis and killing were calculated based on the CFUs in the culture plates. In brief, bacterial CFU at 1 hr post-bacterial stimulation was calculated as CFU/ul (phagocytosis). Bacterial CFU at 2 and 12 hrs post-bacterial stimulation was compared with that at 1 hr to show the percentage of viable bacteria of phagocytosed bacteria.

### *In vivo* depletion of neutrophils

To partially deplete neutrophils *in vivo*, mice were injected i.p with 100μg of anti-Ly6G mAb (clone 1A8 purchased from UCSF Monoclonal Antibody Core, San Fracisco) or 200μg of isotype control mAb (clone 2A3, BioXCell, West Lebanon, NH) at 1 hr post-bacterial challenge. **Lung histopathology**

At selected time points post bacterial infection, lungs were harvested and inflated with 10% formalin for 96 hrs. Images of representative micrographs were taken under a Leica DMRA microscope (Leica Microsystems, Wetzlar, Germany) by using Openlab software version 5.0.2 (Improvi-sion, Coventry, UK).

### Quantification and statistical analysis

Statistical parameters including the exact value of n and statistical significance are reported in Figures and Figure Legends. A p value of < 0.05 was considered significant (*p < 0.05, **p < 0.01, ***p < 0.001, ****p < 0.0001). A two-tailed Student t test was performed for pairwise comparisons and a 2way ANOVA for multiple comparisons. All analyses were performed by using GraphPad Prism software (Version 6, GraphPad Software, La Jolla, CA).

## References

1. Moliva, J. I., J. Turner, and J. B. Torrelles. 2017. Immune responses to bacillus Calmette-Guérin vaccination: Why do they fail to protect against mycobacterium tuberculosis? Front. Immunol..

2. Netea, M. G., L. A. B. Joosten, E. Latz, K. H. G. Mills, G. Natoli, H. G. Stunnenberg, L. A. J. O’Neill, and R. J. Xavier. 2016. Trained immunity: A program of innate immune memory in health and disease. Science (80-.). 352: 427.

3. Xing, Z., S. Afkhami, J. Bavananthasivam, D. K. Fritz, M. R. D’Agostino, M. Vaseghi-Shanjani, Y. Yao, and M. Jeyanathan. 2020. Innate immune memory of tissue-resident macrophages and trained innate immunity: Re-vamping vaccine concept and strategies. J. Leukoc. Biol. 108: 825–834.

4. Pasco, S. T., and J. Anguita. 2020. Lessons from Bacillus Calmette-Guérin: Harnessing Trained Immunity for Vaccine Development. Cells 9.

5. Netea, M. G., L. A. B. Joosten, E. Latz, K. H. G. Mills, G. Natoli, H. G. Stunnenberg, L. A. J. O’Neill, and R. J. Xavier. 2016. Trained immunity: A program of innate immune memory in health and disease. Science (80-.). 352: 427.

6. Xing, Z., S. Afkhami, J. Bavananthasivam, D. Fritz, M. D’Agostino, M. Vaseghi-Shanjani, Y. Yao, and M. Jeyanathan. 2020. Innate immune memory of lung tissue-resident macrophages and trained innate immunity: Re-vamping vaccine concept and strategies. J. Leukoc. Biol..

7. Jensen, K. J., N. Larsen, S. B. Sørensen, A. Andersen, H. B. Eriksen, I. Monteiro, D. Hougaard, P. Aaby, M. G. Netea, K. L. Flanagan, and C. S. Benn. 2015. Heterologous immunological effects of early BCG vaccination in low-birth-weight infants in guinea-bissau: A randomized-controlled trial. J. Infect. Dis. 211: 956–967.

8. Garly, M. L., C. L. Martins, C. Balé, M. A. Baldé, K. L. Hedegaard, P. Gustafson, I. M. Lisse, H. C. Whittle, and P. Aaby. 2003. BCG scar and positive tuberculin reaction associated with reduced child mortality in West Africa: A non-specific beneficial effect of BCG? Vaccine 21: 2782–2790.

9. Stensballe, L. G., E. Nante, I. P. Jensen, P. E. Kofoed, A. Poulsen, H. Jensen, M. Newport, A. Marchant, and P. Aaby. 2005. Acute lower respiratory tract infections and respiratory syncytial virus in infants in Guinea-Bissau: A beneficial effect of BCG vaccination for girls: Community based case-control study. Vaccine 23: 1251–1257.

10. De Castro, M. J., J. Pardo-Seco, and F. Martinón-Torres. 2015. Nonspecific (heterologous) protection of neonatal BCG vaccination against hospitalization due to respiratory infection and sepsis. Clin. Infect. Dis..

11. Walk, J., L. C. J. de Bree, W. Graumans, R. Stoter, G. J. van Gemert, M. van de Vegte-Bolmer, K. Teelen, C. C. Hermsen, R. J. W. Arts, M. C. Behet, F. Keramati, S. J. C. F. M. Moorlag, A. S. P. Yang, R. van Crevel, P. Aaby, Q. de Mast, A. J. A. M. van der Ven, C. Stabell Benn, M. G. Netea, and R. W. Sauerwein. 2019. Outcomes of controlled human malaria infection after BCG vaccination. Nat. Commun. 10.

12. Arts, R. J. W., S. J. C. F. M. Moorlag, B. Novakovic, Y. Li, S. Y. Wang, M. Oosting, V. Kumar, R. J. Xavier, C. Wijmenga, L. A. B. Joosten, C. B. E. M. Reusken, C. S. Benn, P. Aaby, M. P. Koopmans, H. G. Stunnenberg, R. van Crevel, and M. G. Netea. 2018. BCG Vaccination Protects against Experimental Viral Infection in Humans through the Induction of Cytokines Associated with Trained Immunity. Cell Host Microbe 23: 89–100.e5.

13. Giamarellos-Bourboulis, E. J., M. Tsilika, S. Moorlag, N. Antonakos, A. Kotsaki, J. Domínguez-Andrés, E. Kyriazopoulou, T. Gkavogianni, M. E. Adami, G. Damoraki, P. Koufargyris, A. Karageorgos, A. Bolanou, H. Koenen, R. van Crevel, D. I. Droggiti, G. Renieris, A. Papadopoulos, and M. G. Netea. 2020. Activate: Randomized Clinical Trial of BCG Vaccination against Infection in the Elderly. Cell 183: 315–323.e9.

14. Faustman, D. L., A. Lee, E. R. Hostetter, A. Aristarkhova, C. Nathan, G. F. Shpilsky, L. Tran, G. Wolfe, H. Takahashi, H. F. Dias, J. Braley, H. Zheng, D. A. Schoenfeld, and W. M. Kühtreiber. 2022. Multiple BCG vaccinations for prevention of COVID-19 and other infectious diseases in Type 1 diabetes. Cell Reports Med. 100728.

15. Moorlag, S. J. C. F. M., Y. A. Rodriguez-Rosales, J. Gillard, S. Fanucchi, K. Theunissen, B. Novakovic, C. M. de Bont, Y. Negishi, E. T. Fok, L. Kalafati, P. Verginis, V. P. Mourits, V. A. C. M. Koeken, L. C. J. de Bree, G. J. M. Pruijn, C. Fenwick, R. van Crevel, L. A. B. Joosten, I. Joosten, H. Koenen, M. M. Mhlanga, D. A. Diavatopoulos, T. Chavakis, and M. G. Netea. 2020. BCG Vaccination Induces Long-Term Functional Reprogramming of Human Neutrophils. Cell Rep. 33: 108387.

16. Kleinnijenhuis, J., J. Quintin, F. Preijers, L. A. B. Joosten, C. Jacobs, R. J. Xavier, J. W. M. van der Meer, R. van Crevel, and M. G. Netea. 2014. BCG-induced trained immunity in NK cells: Role for non-specific protection to infection. Clin. Immunol. 155: 213–219.

17. Kleinnijenhuis, J., J. Quintin, F. Preijers, L. A. B. Joosten, D. C. Ifrim, S. Saeed, C. Jacobs, J. Van Loenhout, D. De Jong, S. Hendrik, R. J. Xavier, J. W. M. Van Der Meer, R. Van Crevel, and M. G. Netea. 2012. Bacille Calmette-Guérin induces NOD2-dependent nonspecific protection from reinfection via epigenetic reprogramming of monocytes. Proc. Natl. Acad. Sci. U. S. A. 109: 17537–17542.

18. Soto, J. A., N. M. S. Gálvez, C. A. Andrade, M. A. Ramírez, C. A. Riedel, A. M. Kalergis, and S. M. Bueno. 2022. BCG vaccination induces cross-protective immunity against pathogenic microorganisms. Trends Immunol. 43: 322–335.

19. Kaufmann, E., J. Sanz, J. L. Dunn, N. Khan, L. E. Mendonça, A. Pacis, F. Tzelepis, E. Pernet, A. Dumaine, J. C. Grenier, F. Mailhot-Léonard, E. Ahmed, J. Belle, R. Besla, B. Mazer, I. L. King, A. Nijnik, C. S. Robbins, L. B. Barreiro, and M. Divangahi. 2018. BCG Educates Hematopoietic Stem Cells to Generate Protective Innate Immunity against Tuberculosis. Cell 172: 176–190.e19.

20. Cirovic, B., L. C. J. de Bree, L. Groh, B. A. Blok, J. Chan, W. J. F. M. van der Velden, M. E. J. Bremmers, R. van Crevel, K. Händler, S. Picelli, J. Schulte-Schrepping, K. Klee, M. Oosting, V. A. C. M. Koeken, J. van Ingen, Y. Li, C. S. Benn, J. L. Schultze, L. A. B. Joosten, N. Curtis, M. G. Netea, and A. Schlitzer. 2020. BCG Vaccination in Humans Elicits Trained Immunity via the Hematopoietic Progenitor Compartment. Cell Host Microbe 28: 322–334.e5.

21. Kaufmann, E., J. Sanz, J. L. Dunn, N. Khan, L. E. Mendonça, A. Pacis, F. Tzelepis, E. Pernet, A. Dumaine, J. C. Grenier, F. Mailhot-Léonard, E. Ahmed, J. Belle, R. Besla, B. Mazer, I. L. King, A. Nijnik, C. S. Robbins, L. B. Barreiro, and M. Divangahi. 2018. BCG Educates Hematopoietic Stem Cells to Generate Protective Innate Immunity against Tuberculosis. Cell 172: 176–190.e19.

22. Mata, E., R. Tarancon, C. Guerrero, E. Moreo, F. Moreau, S. Uranga, A. B. Gomez, D. Marinova, M. Domenech, F. Gonzalez-Camacho, M. Monzon, J. Badiola, J. Dominguez-Andres, J. Yuste, A. Anel, A. Peixoto, C. Martin, and N. Aguilo. 2021. Pulmonary BCG induces lung-resident macrophage activation and confers long-term protection against tuberculosis. Sci. Immunol. 6: 1–14.

23. Vierboom, M. P. M., K. Dijkman, C. C. Sombroek, S. O. Hofman, C. Boot, R. A. W. Vervenne, K. G. Haanstra, M. van der Sande, L. van Emst, J. Domínguez-Andrés, S. J. C. F. M. Moorlag, C. H. M. Kocken, J. Thole, E. Rodríguez, E. Puentes, J. H. A. Martens, R. van Crevel, M. G. Netea, N. Aguilo, C. Martin, and F. A. W. Verreck. 2021. Stronger induction of trained immunity by mucosal BCG or MTBVAC vaccination compared to standard intradermal vaccination. Cell Reports Med. 2.

24. Kaufmann, E., N. Khan, K. A. Tran, A. Ulndreaj, E. Pernet, G. Fontes, A. Lupien, P. Desmeules, F. McIntosh, A. Abow, S. J. C. F. M. Moorlag, P. Debisarun, K. Mossman, A. Banerjee, D. Karo-Atar, M. Sadeghi, S. Mubareka, D. C. Vinh, I. L. King, C. S. Robbins, M. A. Behr, M. G. Netea, P. Joubert, and M. Divangahi. 2022. BCG vaccination provides protection against IAV but not SARS-CoV-2. Cell Rep. 38.

25. Yao, Y., M. Jeyanathan, S. Haddadi, N. G. Barra, M. Vaseghi-Shanjani, D. Damjanovic, R. Lai, S. Afkhami, Y. Chen, A. Dvorkin-Gheva, C. S. Robbins, J. D. Schertzer, and Z. Xing. 2018. Induction of Autonomous Memory Alveolar Macrophages Requires T Cell Help and Is Critical to Trained Immunity. Cell 175: 1634–1650.e17.

26. Knapp, S., J. C. Leemans, S. Florquin, J. Branger, N. A. Maris, J. Pater, N. Van Rooijen, and T. Van der Poll. 2003. Alveolar macrophages have a protective antiinflammatory role during murine pneumococcal pneumonia. Am. J. Respir. Crit. Care Med. 167: 171–179.

27. Yao, Y., M. Jeyanathan, S. Haddadi, N. G. Barra, M. Vaseghi-Shanjani, D. Damjanovic, R. Lai, S. Afkhami, Y. Chen, A. Dvorkin-Gheva, C. S. Robbins, J. D. Schertzer, and Z. Xing. 2018. Induction of Autonomous Memory Alveolar Macrophages Requires T Cell Help and Is Critical to Trained Immunity. Cell 175: 1634–1650.e17.

28. Mai, D., A. Jahn, T. Murray, M. Morikubo, J. Nemeth, K. Urdahl, A. H. Diercks, A. Aderem, and A. C. Rothchild. 2022. Mycobacterial exposure remodels alveolar macrophages and the early innate response to &lt;em&gt;Mycobacterium tuberculosis&lt;/em&gt; infection. bioRxiv 2022.09.19.507309.

29. Delahaye, J. L., B. H. Gern, S. B. Cohen, C. R. Plumlee, S. Shafiani, M. Y. Gerner, and K. B. Urdahl. 2019. Cutting Edge: Bacillus Calmette–Guérin–Induced T Cells Shape Mycobacterium tuberculosis Infection before Reducing the Bacterial Burden. J. Immunol. 203: 807–812.

30. Vierboom, M. P. M., K. Dijkman, C. C. Sombroek, S. O. Hofman, C. Boot, R. A. W. Vervenne, K. G. Haanstra, M. van der Sande, L. van Emst, J. Domínguez-Andrés, S. J. C. F. M. Moorlag, C. H. M. Kocken, J. Thole, E. Rodríguez, E. Puentes, J. H. A. Martens, R. van Crevel, M. G. Netea, N. Aguilo, C. Martin, and F. A. W. Verreck. 2021. Stronger induction of trained immunity by mucosal BCG or MTBVAC vaccination compared to standard intradermal vaccination. Cell Reports Med. 2.

31. Kleinnijenhuis, J., J. Quintin, F. Preijers, L. A. B. Joosten, D. C. Ifrim, S. Saeed, C. Jacobs, J. Van Loenhout, D. De Jong, S. Hendrik, R. J. Xavier, J. W. M. Van Der Meer, R. Van Crevel, and M. G. Netea. 2012. Bacille Calmette-Guérin induces NOD2-dependent nonspecific protection from reinfection via epigenetic reprogramming of monocytes. Proc. Natl. Acad. Sci. U. S. A. 109: 17537–17542.

32. Cirovic, B., L. C. J. de Bree, L. Groh, B. A. Blok, J. Chan, W. J. F. M. van der Velden, M. E. J. Bremmers, R. van Crevel, K. Händler, S. Picelli, J. Schulte-Schrepping, K. Klee, M. Oosting, V. A. C. M. Koeken, J. van Ingen, Y. Li, C. S. Benn, J. L. Schultze, L. A. B. Joosten, N. Curtis, M. G. Netea, and A. Schlitzer. 2020. BCG Vaccination in Humans Elicits Trained Immunity via the Hematopoietic Progenitor Compartment. Cell Host Microbe.

33. Taut, K., C. Winter, D. E. Briles, J. C. Paton, J. W. Christman, R. Maus, R. Baumann, T. Welte, and U. A. Maus. 2008. Macrophage turnover kinetics in the lungs of mice infected with Streptococcus pneumoniae. Am. J. Respir. Cell Mol. Biol. 38: 105–113.

34. Santosuosso, M., S. McCormick, X. Zhang, A. Zganiacz, and Z. Xing. 2006. Intranasal boosting with an adenovirus-vectored vaccine markedly enhances protection by parenteral Mycobacterium bovis BCG immunization against pulmonary tuberculosis. Infect. Immun..

35. Jeyanathan, M., S. Afkhami, A. Khera, T. Mandur, D. Damjanovic, Y. Yao, R. Lai, S. Haddadi, A. Dvorkin-Gheva, M. Jordana, S. L. Kunkel, and Z. Xing. 2017. CXCR3 Signaling Is Required for Restricted Homing of Parenteral Tuberculosis Vaccine–Induced T Cells to Both the Lung Parenchyma and Airway. J. Immunol. 199: 2555–2569.

